# Inhibition of Phosphodiesterase 12 results in Antiviral Activity against several RNA Viruses including SARS-CoV-2

**DOI:** 10.1101/2022.09.23.509178

**Authors:** Mark Thursz, Fouzia Sadiq, Julia A. Tree, Peter Karayiannis, David W. C. Beasley, Wanwissa Dejnirattisai, Juthathip Mongkolsapaya, Gavin Screaton, Matthew Wand, Michael J. Elmore, Miles W. Carroll, Ian Matthews, Howard Thomas

## Abstract

Phosphodiesterase 12 (PDE12) is a negative regulator of the type 1 interferon (IFN) response and here we show that PDE12 inhibitors (lead compounds 63 and 17) are associated with increased RNAseL activity, are well tolerated at the therapeutic range and inhibit, both *in vitro* and *in vivo*, the replication of several RNA viruses including hepatitis C virus (HCV), dengue virus (DENV), West Nile Virus (WNV) and SARS-CoV-2.

## Introduction

The host IFN-induced antiviral response requires the activation of several effector pathways including that of 2’,5’-oligoadenylate synthetase (OAS) – Ribonuclease L (RNAseL). OAS transcription is induced by IFN and, when the enzyme is activated by double-stranded RNA (dsRNA), a replicative intermediate of RNA viruses, it generates a unique signalling intermediate, 2’,5’-oligoadenylate(2’5’OA) using ATP as a substrate. 2’*5*’OA binds to the RNAseL, resulting in dimerization and activation. Activated RNAseL hydrolyses RNA species fairly indiscriminately resulting in the termination of viral replication but may also result in infected cell death via apoptosis due to cleavage of cellular RNAs (1). Consequently, the OAS-RNAseL pathway requires tight regulation which is provided by the enzyme phosphodiesterase 12 (PDE12) which rapidly hydrolyses 2’5’OA to terminate RNAseL activity (2).

The OAS-RNAseL pathway is thought to be an effective innate immune response against a wide range of human and animal viruses (3). Gene association studies have implicated this pathway as a determinant of outcome for hepatitis C virus (HCV) infection (4) and more recently a genome wide association study identified polymorphisms in the OAS1, OAS2, OAS3 cluster on chromosome 19 correlated with the severity of COVID-19 lung disease (5).

We speculated that inhibition of PDE12 to manipulate the activity of RNAseL could be used therapeutically to inhibit viral replication. The requirement for dsRNA to activate OAS should provide specificity for RNAseL activation to be restricted to viral infected cells. Here we show that inhibition of PDE12 with a small molecule inhibitor, is well tolerated in rodents and results in increased levels of intracellular RNAseL activity in response to virus infection (EMCV). In addition, we show that these PDE12 inhibitors have antiviral activity *in-vitro* against clinically important RNA viruses including DENV, WNV, HCV, and SARS-CoV-2. In early studies in mice infected with WNV, we show reduced viraemia and mortality.

## Materials and Methods

### Cell Culture

Human Huh7 hepatoma, Huh7.5 HCV permissive hepatocyte, Human Embryonic kidney 293T and the mutant U5A (non-responsive to type 1 IFNs) cell lines were cultured at 37°C in growth media of Dulbecco’s Modified Eagle Medium (DMEM; Invitrogen Ltd, Paisley, UK) supplemented with 10% Foetal Bovine Serum (FBS) and penicillin/streptomycin in a humidified environment of 5% CO_2_.

Human epithelial (HEp-2) and Madin Darby Bovine Kidney (MDBK) cells were grown in RPMI and MEM supplemented with 2% FBS media, respectively. Incubations with virus were performed in either serum free or media supplemented with 2% FBS without penicillin/streptomycin.

VeroE6 cells (ECACC 85020206) were grown in MEM supplemented with 10% FBS, L-glutamine, non-essential amino acids and 25mM HEPES (8).

A549 and A549-ACE2 cells were grown in MEM supplemented with 10% FBS, penicillin/streptomycin, and L-glutamine. Incubations with virus were performed in media supplemented with 2% FBS, penicillin/streptomycin, and L-glutamine.

### Most antiviral assays

were set up in 96 well cell culture plates and seeded with 2×10^4^ cells per well. For WNV and SARS-CoV-2, 12-well plates were seeded with 4×10^5^ A549 or A549-ACE2 cells. Murine encephalomyocarditis virus (EMCV) was grown in our laboratory while DENV serotypes, WNV and Respiratory Syncytial Virus (RSV), were grown in the laboratories of the respective collaborators.

Antiviral assays were employed to determine the ability of PDE12 inhibitors to potentiate interferon (IFN) activity on viral titres, unless otherwise noted. Briefly during EMCV assay, Huh7 hepatoma cells were stimulated overnight with a fixed dose of IFNα (1600pg/ml) in growth medium, prior to a challenge with EMCV at a multiplicity of infection (MOI) of 0.05 for an hour. EMCV was removed and cells were incubated in growth medium supplemented with or without PDE12 inhibitors for 24 hours at 37°C. At the end of incubation, cells were stained with crystal violet and absorbance was read at 540nm. Absorbance is directly proportional to cell viability.

### Realtime quantitative PCR

To quantify EMCV viral load, Huh7 cells were stimulated with IFNα and challenged with EMCV as described under antiviral assay. After incubating Huh7 cells in growth medium supplemented with PDE12 inhibitors for 20 hours, samples of supernatant were removed for RNA extraction. Tri Reagent (SIGMA-Aldrich Ltd, Dorset, UK) was used to isolate RNA following the manufacturer’s guidelines. RNA (~1μg) was reversed transcribed to cDNA using the RETROscript^®^ kit (Ambion Applied Biosystems, UK). Quantitative RT-PCR was performed using the SYBR Green detection system (Qiagen) according to the manufacturer’s instructions in an iCycler (Bio-Rad Laboratories Ltd., UK). The pair of primers used for amplification of a 116bp fragment of the EMCV genome was: Forward primer 5’-TAC CGG GGA TCA TTG GTT TA-3’, Reverse primer 5’-GTC TCG ACT AGT GGG CTT GC-3’.

### Taqman assays

For PDE12 and RNaseL gene expression studies quantitative polymerase chain reaction (qPCR) analysis was performed using TaqMan Gene Expression Assays (ThermoFisher, UK). Data were normalized to the endogenous housekeeping gene GADPH and fold change differences in expression relative to control cells were calculated using the Comparative ddCT method.

### Cell Toxicity assay

Huh7 cells were seeded in a 96-well culture plate at a density of 2×10^4^ cells per well and incubated for 24 hours with PDE12 inhibitor at various concentrations as indicated in figure legends. At the end of incubations, the medium was removed and cells were stained with Crystal Violet. Absorbance was read at 540nm.

### Transfection of T293 cells

T293 cells were transfected with plasmid DNA (pGIPZ shPDE12) using Lipofectamine 2000 reagent (Invitrogen) following the manufacturer’s instructions. The transfected cells were incubated at 37°C for 24 or 48 hours in a humidified environment of 5% CO_2_. Cells were then stimulated with IFN-α and challenged with EMCV as described under the appropriate Figure legends.

### Mouse pharmacokinetics (pK) and rat toxicity analyses

Both pK and *in vivo* toxicity analyses were outsourced to Cyprotex (Cheshire, UK) and Covance Laboratories Ltd (Harrogate, North Yorkshire), respectively. For pK analyses, mice (n=3) were administered either intravenously (<u>IV</u>) or intraperitoneally (<u>IP</u>) or orally with CO-63 at 0.14, 0.28 or 0.29 mg/kg, respectively. Plasma samples were collected at various time points for up to 8 hours and CO-63 was analysed. For toxicity analyses, rats were administered with CO-63 at 25, 50, 75 or 100mg/kg and acute minimum lethal intravenous dose of the compound was determined.

### Mouse studies of anti-West Nile virus efficacy

Mouse studies <u>to assess antiviral efficacy against neuroinvasive WNV disease in a mouse model</u> were undertaken at the University of Texas Medical Branch (UTMB). <u>Studies</u> <u>were</u> carried out in strict accordance with the recommendations in the Guide for the Care and Use of Laboratory Animals of the National Institutes of Health under a. protocol approved by the UTMB Institutional Animal Care and Use Committee. <u>UTMB is a registered Research Facility under the Animal Welfare Act,. has a current assurance on file with the Office of Laboratory Animal Welfare (OLAW), and is AAALAC (Association for the Assessment and Accreditation of Laboratory and Care International) accredited. Groups of mice (n=10/group) were challenged with a lethal dose of WNV (100 pfu, New York 1999 strain) then treated via the IP route with 50 mg/kg CO-63 or sterile PBS at 4, 8, 12, 16 and 24 hours post-challenge. Mice were monitored up to 3 times daily (morning, afternoon and night during peak of disease) and scored for body weight changes and other clinical signs of WNV disease (e.g. ruffled fur, hunching, lethargy, limb paralysis) for 21 days. Animals meeting pre-determined criteria were humanely euthanized. To measure viremia levels, blood samples were collected from mice at day 3 or 4 post-challenge via the retro-orbital route, under inhalational anesthesia, and virus titers in serum determined by standard plaque assay on Vero cells.</u>

<u>No patient samples were used in this study and therefore Ethical approval or informed consent were not required</u>.

## Results

### PDE-12 activity assay

In order to perform high throughput screening an assay was developed to measure PDE12 activity using a 2’5’ linked oligoadenylate probe with a fluorophore at the 2’ end and a quencher at the 5’ end 2’5’-OA, (dR)FAM-A_2p5_A_2p5_A_2p5_A_2p5_A_2p5_A-DDQ-1 (Eurogentec Ltd, Southampton, UK) (Figure 1A). Reaction mixtures contained cell lysate from HepG2 cells which express high levels of PDE12 and the substrate at a concentration of 6μM. Nerve growth factor (NGF) inhibits PDE12 activity so, in the absence of a specific PDE12 inhibitor, lysate from NGF-treated HepG2 cells was used as a negative control and untreated HepG2 lysate as a positive control. Fluorescence (excitation at 485 nm; emission at 520 nm) was measured twice using the BMG LabTech Flourostar at Time zero and 120 min of the assay. Figure 1B illustrates the rapid dynamic of PDE12 activity and the response to NGF treatment.

**Figure 1A.**
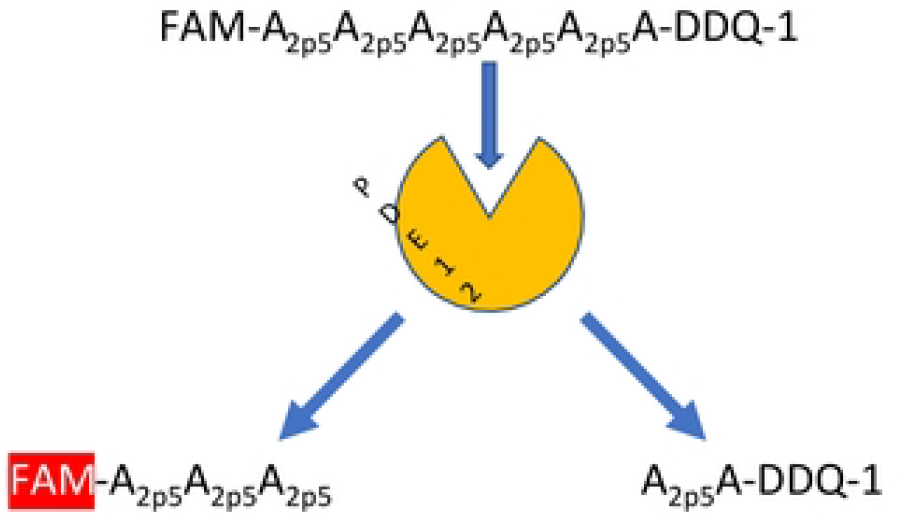
PDE-12 activity assay. The PDE12 Activity Assay employed to perform high throughput screening (HTS) for inhibitors of PDE12 (defined as ≥50% inhibition of PDE12 activity), from a library of 18000 small molecules, resulting in 6 hits.

**Figure 1B.**
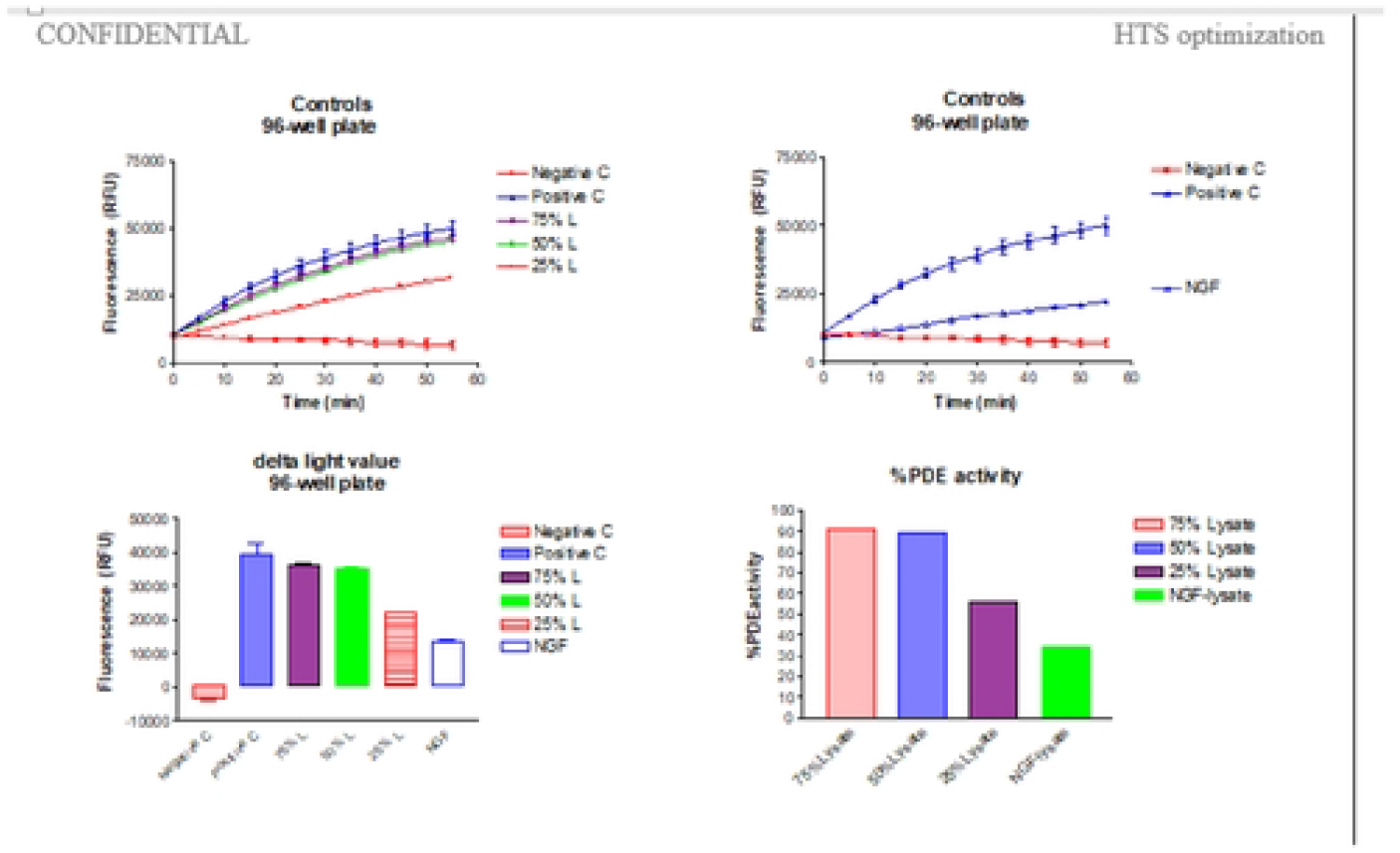
Dynamics of PDE12 activity. Rapid dynamic of PDE12 activity and the response to NGF treatment. C, control; L, lysate; NGF, nerve growth factor.

### Screening and Lead Development

Primary screening of ~18,000 compounds at a concentration of 10μM resulted in the identification of 6 hits where PDE12 activity was reduced by >50% of the positive control. To increase the repertoire of the lead molecule, 160 compounds based on structural similarities, for focussed screening were purchased from ChemBridge, ChemDiv, ChemBlock and InterBioScience. Novel small molecules (N=174) were synthesized at ChemOvation (Horsham, West Sussex, UK). All molecules were dissolved in DMSO and diluted in assay buffer or cell culture media prior to use. Lead compounds generated through this process were tested against a range of phosphodiesterases to determine specificity. Apart from low level inhibition of PDE7, no significant ‘off-target’ activity was observed (Supplementary table 1).

### Enhanced antiviral activity against Encephalomyocarditis Virus (EMCV)

Antiviral assays using EMCV, a picornavirus, is widely used to assess interferon effector pathways. In the Huh-7 hepatoma cell line infected with EMCV at a MOI of 0.05, the IFN α2 dose response curve was shifted left by 1.5 log in potency: IC_50_ of IFN α2 plus vehicle was 3,536pg/ml and the IC_50_ of IFN α2 plus CO-63 was 270.4 pg/ml (Figure 1C)

**Figure 1C.** PDE12 enhances the antiviral effect of interferon. PDE12 inhibitors enhance the antiviral effect of type 1 IFN. EMCV in the supernatants from antiviral assays was used as inoculum and incubated with A549 cells for an hour. These were then removed and cells were overlaid with growth medium supplemented with carboxy methyl cellulose (CMC) for 24 hours. Cells were stained with methyl violet and lytic plaques were counted.

### Ribosomal RNA cleavage

Activation of RNAseL will result in cleavage of cellular RNA species in addition to viral RNA. To demonstrate that the enhanced antiviral effect of PDE12 inhibition is due to increased RNAseL activity, total cellular RNA was extracted from cells infected with EMCV and treated with IFN α2 in the presence or absence of CO-63 at a concentration of 20μM. Ribosomal RNA cleavage fragments appeared at earlier time-points and w<u>ere</u> more intense in cells treated with the PDE12 inhibitors (Figure 1D).

**Figure 1D.**
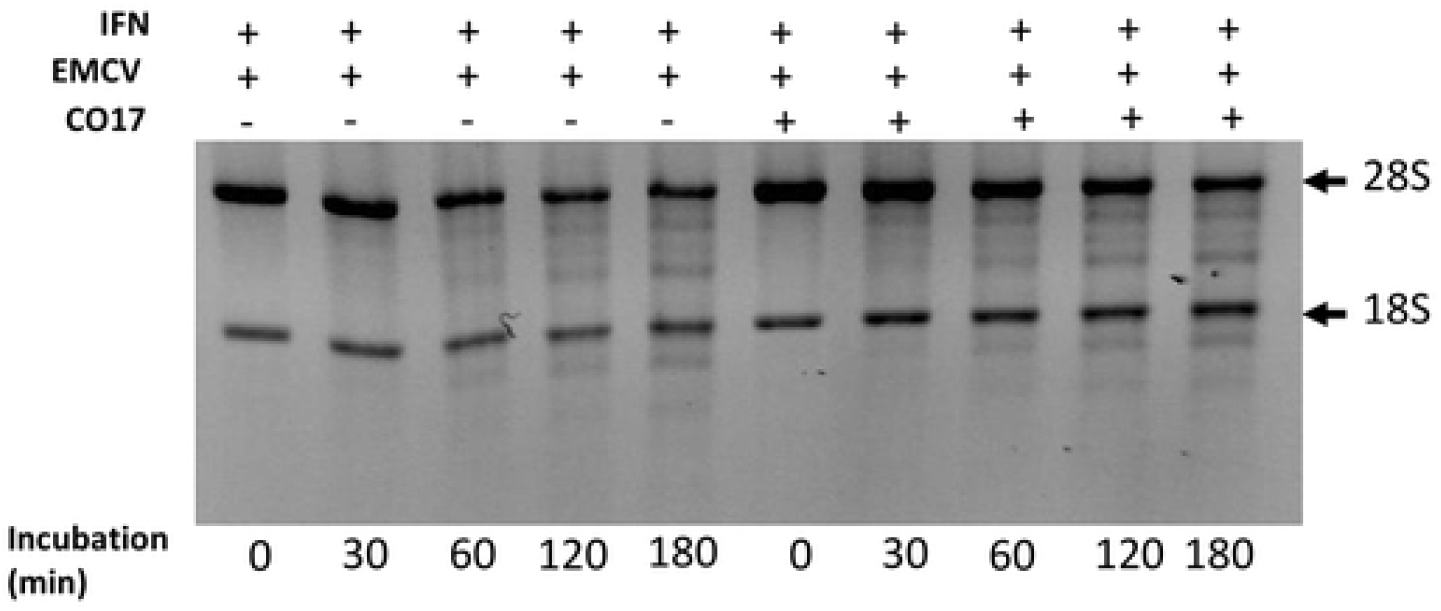
PDE12 inhibition increases ribosomal RNA cleavage. Activation of RNAseL resulted in cleavage of cellular RNA species in addition to viral RNA. Total cellular RNA was extracted from cells infected with EMCV and showed rRNA (18s and 28s) cleavage fragments appearing at earlier timepoints and being more intense in cells treated with IFN a2 in the presence of CO-63 at a concentration of 20mM. Huh-7 cell pellets were gently lysed in 1.5 pellet volume of lysis buffer (10mM Hepes, pH 7.5, 90mM KCl, 1mM Magnesium acetate, 0.5% (v/v) Triton X-100, 2mM fresh mercaptoethanol and 100μg/ml fresh Leupeptin) on ice for 5 min. Crude lysate was centrifuged for 10,000g at 4°C for 10 min and the supernatant was transferred to a clean tube. The rRNA cleavage reaction was set up in 10X cleavage buffer (100mM Hepes, pH 7.5, 1M KCl, 50mM Magnesium acetate, 10mM ATP, 0.14 M 2 mercaptoethanol). Cleared lysate (20μg protein) was incubated in 1X cleavage buffer for 0, 0.5, 1, 2 and 3 hours at 30°C. RNA was isolated from incubated lysate using the RNeasy kit (Qiagen, GmBH Germany) according to the manufacturer’s instructions. RNA was quantified by measuring absorbance at 260nm and nanomolar concentration of RNA was separated on RNA chips and analysed with an Agilent Bioanalyser 2100 (Agilent Technologies).

### In vitro Anti-Viral Activity

The spectrum of *in vitro* antiviral activity was evaluated against a range of viruses including HCV, DENV, WNV and SARS-CoV-2.

Antiviral activity against HCV was tested using Huh7 cells infected with the genotype 2 JFH1 strain of the virus. In Huh7 cells, treatment with CO-63 at concentrations ranging from 20-40μM in the presence of IFN α2 at a concentration of 20pg/ml, suppressed JFH1 markedly (Figure 2A).

**Figure 2A.**
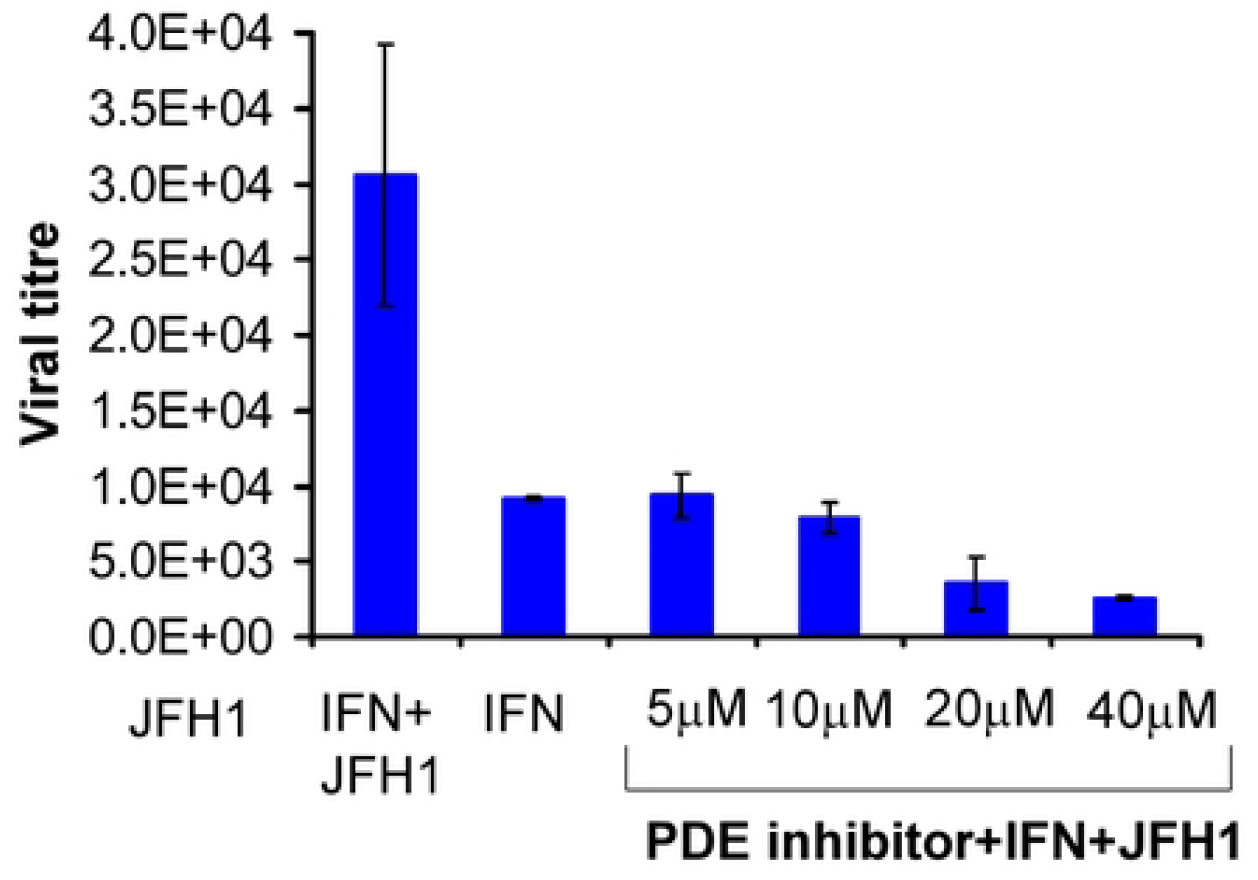
Antiviral activity against the JFH1 strain of HCV. Huh7.5 cells were infected with JFH1 virus (MOI=3) and incubated at 37°C for 48 hours for viral replication. The medium was replaced with media supplemented with IFN α2 (25 IU/ml) for 24 hours. PDE12 inhibitor at indicated concentrations was added and cells were incubated for another 24 hours. Intracellular inhibition of viral replication was determined using the SYBR Green qRT-PCR Kit (Qiagen). Primer sequences employed for amplification of a 131bp product were: Forward 5’-TCTGCGGAACCGGTGAGTA-3’, Reverse 5’-TCAGGCAGTACCACAAGGC-3’. HCV gene expression was normalized to house-keeping gene GAPDH (Glyceraldehyde 3-Phosphate dehydrogenase) expression. Primers for GAPDH amplification were: Forward 5’-TCTCTGCTCCTCCTGTTCGAC-3’ and Reverse 5’-CGGATTTGGTCGTATTGGG-3’.

Antiviral activity against DENV was present in hepatocytes and dendritic cells in the absence of added IFN α2 presumably because the virus triggered endogenous IFN production and in the presence of added IFN antiviral activity was enhanced by compounds 63 (p <0.003) and 17 (p <0.03) in both hepatocyte (Huh-7) cells and monocyte derived dendritic cells (MDDM). In these experiments viral replication was quantitated by measuring the number of cells expressing the DENV-NS1 protein detected by flow cytometry (Figure 2B). Cells were incubated with CO-63 or CO-17 at concentrations of 10, 20 or 40 μM in the presence (1 or 10pg/ml) or absence of IFN α2. In the absence of IFN, CO-17 inhibited viral replication at the 40 μM concentration whereas in the presence of IFN, CO-17 enhanced the antiviral effect even at concentrations of 10 μM (figure 2B).

**Figure 2B.**
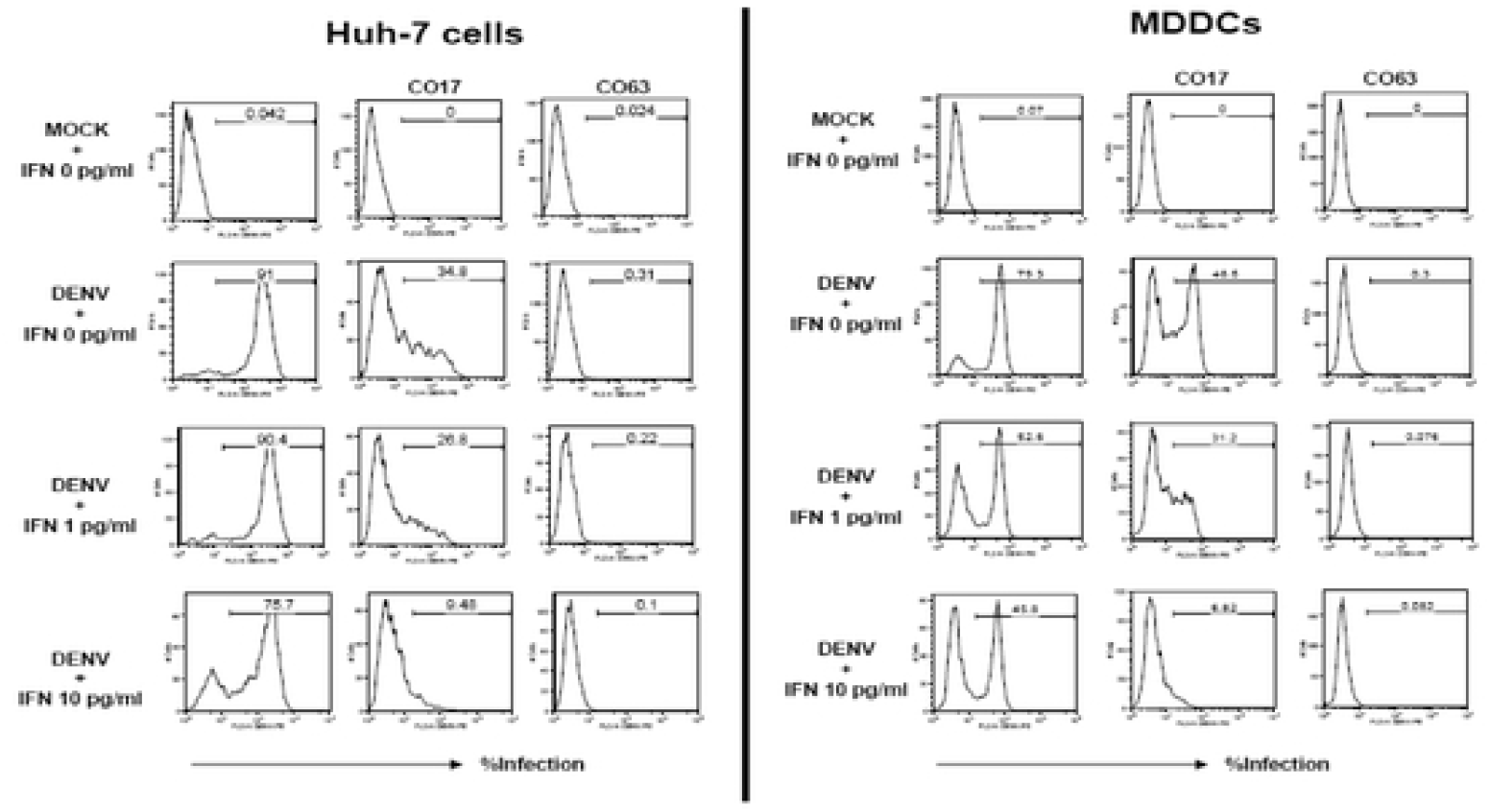
Antiviral activity against DENV. PDE12 inhibitors were investigated in a standard DENV antiviral assay *in vitro*. Antiviral activity against DENV was present in hepatocytes and dendritic cells in the absence of added alpha interferon presumably because the virus triggered endogenous interferon production and in the presence of added interferon antiviral activity was enhanced by CO-63 (p <0.003) and CO-17 (p <0.03) in both hepatocytes (Huh7) cells and monocyte derived dendritic cells (MDDM). In these experiments viral replication was quantitated by measuring the number of cells expressing the DenV-NS1 protein detected by flow cytometry. Cells were incubated with CO-63 or CO-17 at concentrations of 10, 20 or 40 mM in the presence (1 or 10pg/ml) or absence of interferon a2. In the absence of interferon, CO-17 inhibited viral replication at the 40 mM concentration whereas in the presence of interferon, CO-17 enhanced the antiviral effect even at concentrations of 10 mM Initially, Huh7 or Dendritic cells were stimulated with or without IFN α2 at 37°C. After 24 hours, cells were challenged with Dengue virus serotype 2 (16681) for 2 hours. PDE12 inhibitors were included either 2 hours prior to virus challenge or post infection and cells were incubated for another 24 hours. Cells were stained for DENV-NS1 and analysed using flow cytometry. The figure on the right shows that Huh7 cells pre-stimulated with IFN α2 are protected against DENV replication. However, the cells which were not stimulated with IFN α2 allowed DENV replication. Addition of PDE12 inhibitor (CO-17) protected the cells against DENV in a dose dependent fashion. CO-17 at 40μM yielded a 2 log shift towards the left. Stimulation of Huh7 cells with PDE12 inhibitor 2 hours prior to DENV challenge protected the cells against infection even at 10μM, in both monocyte derived dendritic cells and Huh-7 cells using the expression of the DENV-NS1 protein detected by flow cytometry to assess viral replication. Cells were incubated with CO-17 at concentrations of 10, 20 or 40 μM in the presence (1 or 10pg/ml) or absence of IFN α2. In the absence of IFN CO-17 inhibited viral replication at the 40 μM concentration whereas in the presence of IFN CO-17 enhanced the antiviral effect even at concentrations of 10 μM.

Antiviral activity against WNV was assessed in human respiratory carcinoma cells (A549) infected with WNV strain NY99 (7) at an MOI of 1 in the presence of IFN α2 at a concentration of 400 pg/ml. PDE12 inhibition using CO-63 at a concentration of 20 μM resulted in an additional 3 log_10_ reduction in viral RNA at 24 hours and 48 hours compared to IFN alone (Figure 2C).

**Figure 2C.**
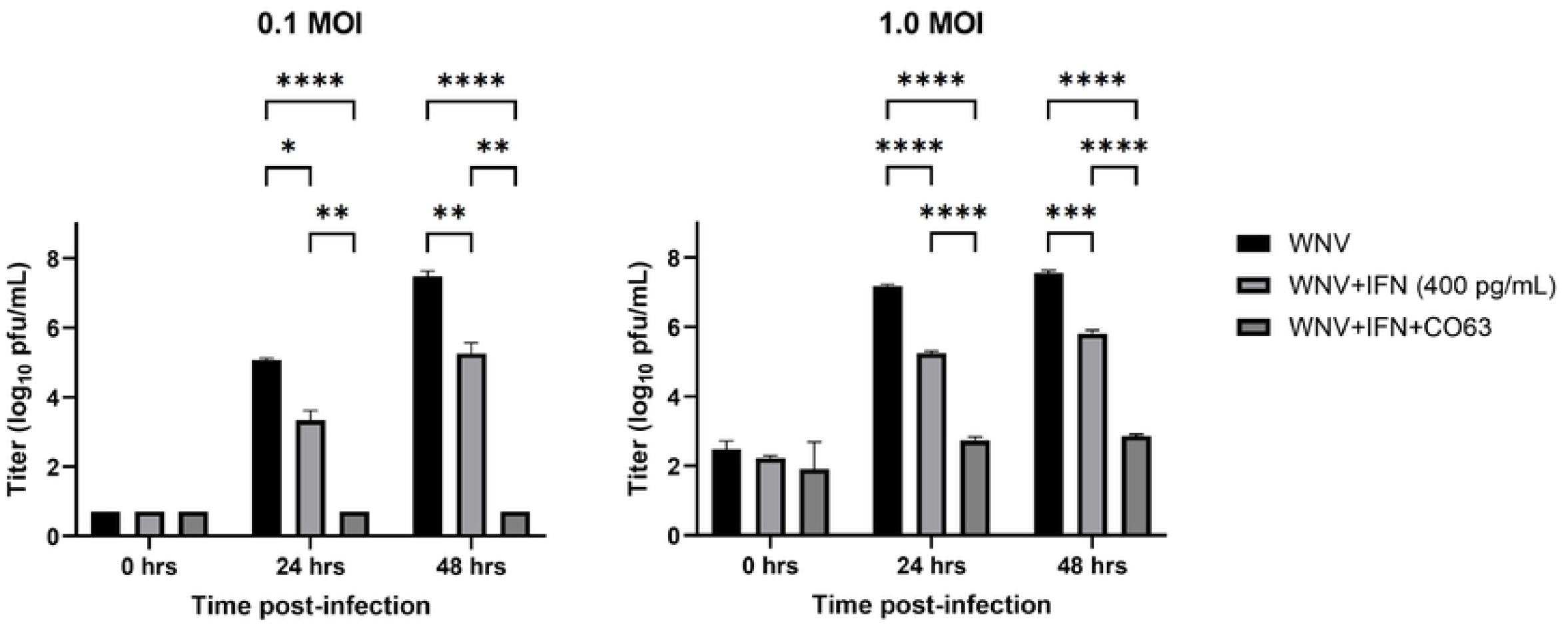
Antiviral activity against WNV. Antiviral activity against WNV was assessed in A549 cells infected with WNV strain NY99 at MOIs of 0.01 and 1 in the presence of IFN α2 at a concentration of 400 pg/ml. PDE12 inhibition using CO-63 at a concentration of 20 μM resulted in an additional 3 log10 reduction in viral RNA at 24 hours and 48 hours compared to IFN alone. Significance values from 2-way ANOVA with Tukey’s multiple comparisons test: * - 0.01 to <0.05,** - 0.001 to <0.01,*** - 0.0001 to <0.001,**** - <0.0001.

Antiviral activity against the SARS-CoV-2 isolate hCoV-19/Australia/VIC01/2020, was evaluated using a microfoci viral-inhibition assay with VeroE6 cells in 96 well plates. Following incubation at 37°C for 24 hours cells and virus were fixed with 8% formalin and an immunostaining protocol (8) was used to visualise the foci. Foci were then counted using an ELISPOT counter (Cellular Technology Ltd) and the data was imported into R-Bioconductor and a mid-point probit analysis was used to determine the IC_50_. The geometric mean IC_50_ for CO-63, in the absence of exogenous IFNα, was 17.11 μg/ml and in the presence of 10 pg/ml of IFNα was 1.28 μg/ml. The geometric mean IC_50_ for CO-17, in the absence of exogenous interferon α, was 10.73 μg/ml and in the presence of 10 pg/ml of IFNα was 0.23 μg/ml (data from experiments conducted at UKHSA, Porton Down, UK).

We confirmed these data using ACE-2 transfected human respiratory carcinoma cells (A549) (kindly provided by Dr. Kent Tseng, UTMB) infected with SARS-CoV-2 isolate USA-WA1/2020. Again, we showed that type 1 IFN produced modest inhibition of viral replication which was substantially enhanced by both PDE12 inhibitors (CO-63 and −17) (Figure 2D).

**Figure 2D:**
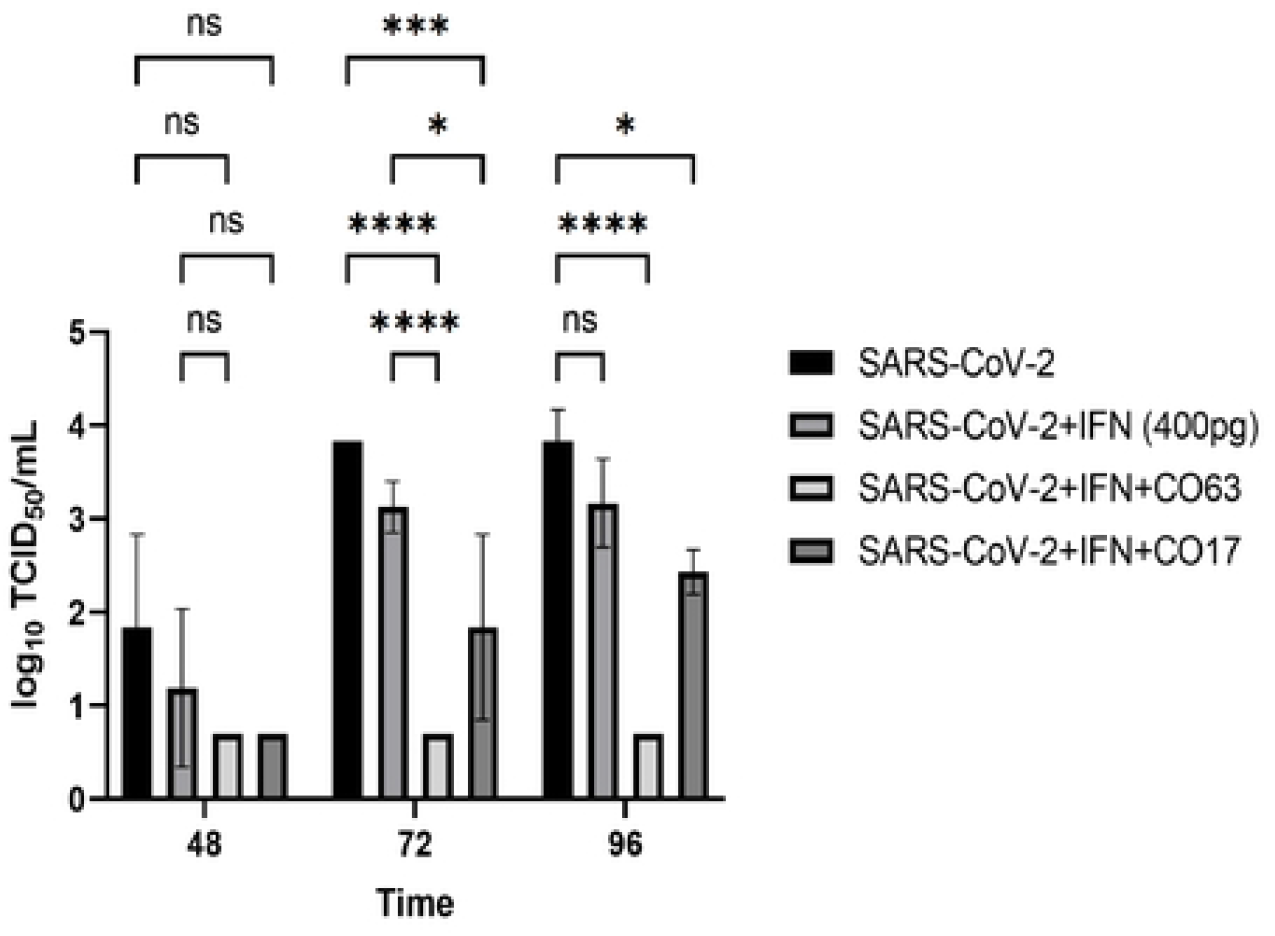
Antiviral activity SARS-CoV-2 (A549 cells expressing ACE2). Antiviral activity against SARS-CoV-2 in A549 cells using a MOI of 1 and measuring titres at the time points and conditions of culture shown. Results are the average of triplicates and statistical significance was determined by two-way ANOVA with Tukey multiple comparisons test in Prism 9 (GraphPad Software, LLC). * - 0.01 to <0.05, ** - 0.001 to <0.01, *** - 0.0001 to <0.001, **** - <0.0001.

### *In-Vivo* Activity

*In-vivo* antiviral activity was tested against WNV in 6–7-week-old female C57BL/6 mice challenged by intraperitoneal injection of a lethal dose of WNV (100pfu); CO-63 (50 mg/kg) or vehicle were given by i.p. injection on days 0, 1 (4, 8, 12, 16, 24 hours). Animals were followed for up to <u>21</u> days, during which the untreated animals lost 13% of their baseline weight compared to 3% in the treated animals, (p<0.05). All animals in the untreated group <u>were euthanized or succumbed to disease</u> -by day 10, whereas 50% of the animals in the treated group survived <u>to the end of the study</u> (Figure 3A).

**Figure 3A:**
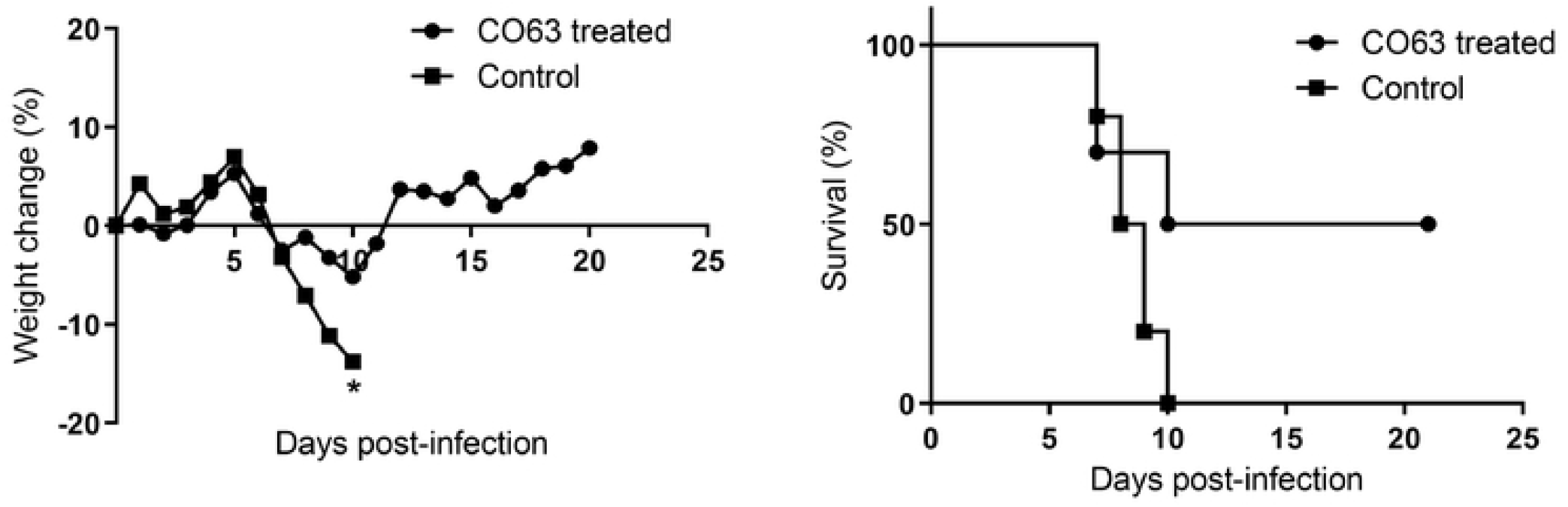
In-Vivo Antiviral Activity against WNV (NY99) in Swiss Webster mice. *In-vivo* antiviral activity was tested against WNV in 6-7 week-old female C57BL/6 mice challenged by intraperitoneal injection of a lethal dose of WNV (100pfu); CO-63 (50 mg/kg) or vehicle were given by i.p. injection on days 0, 1 (4, 8, 12, 16,24 hours). Animals were followed for up to -<u>21</u> days, during which the untreated animals lost 13% of their baseline weight compared to 3% in the treated animal, (p<0.05). All animals in the untreated group <u>were euthanized (n=9) or succumbed to disease (n=1)</u> by day 10, whereas 50% of the animals in the treated group survived. Survival curve statistics: Mantel-Cox test shows statistical significance (p = 0.021) between CO63 treated and control (calculated in Graphpad Prism v9. A. Weight loss and B. survival.

In a separate group of Swiss Webster (SW) 5–6-week-old female mice challenged and treated as above, virological titres measured by plaque assay <u>on serum samples collected</u> on day 3 <u>post-challenge</u>-, showed a significant reduction (p< 0.05) (Figure 3B).

**Figure 3B:**
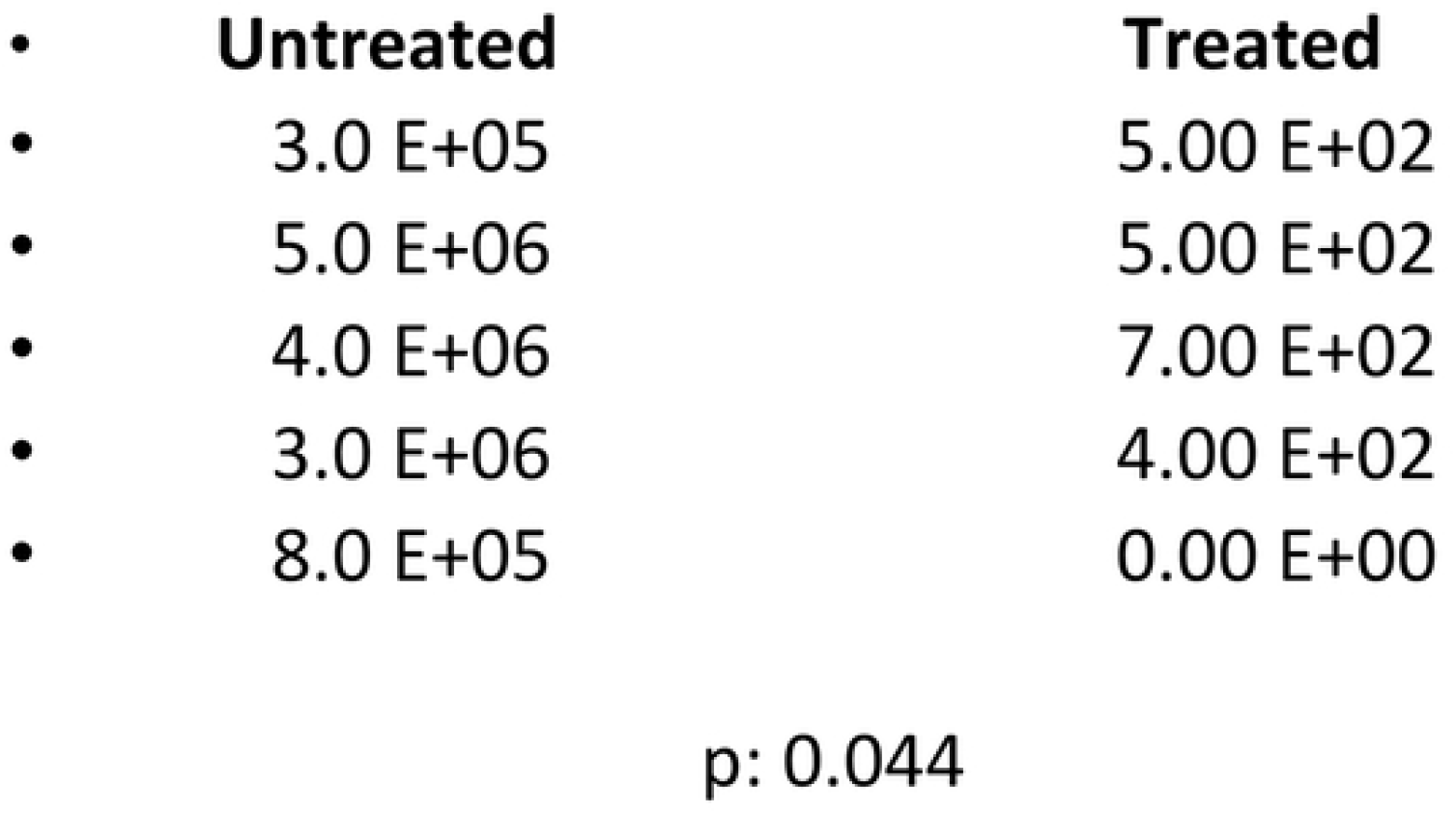
*In-vivo* antiviral activity against WNV (NY99) in Swiss Webster mice: viral titers on day 3. In a separate group of Swiss Webster (SW) 5-6 week-old female mice challenged and treated as in Figure 3A, virological titre was measured by plaque using blood obtained by retro-orbital puncture taken on day 3. A statistically significant difference (p < 0.05) in body-weight change at 10 dpi was apparent using 2-way ANOVA with Sidak test for multiple comparisons.

## Discussion

Our results indicate that PDE12 is readily inhibited by small molecules which results in the amplification of cytoplasmic activity of RNAseL. Furthermore, we have shown that this pathway can be utilised to reduce or prevent viral replication against a range of RNA viruses. Reduction in WNV related mortality in mice provides support for the therapeutic evaluation of these compounds in animal models and potentially, after further toxicity testing, in the human population. In mutant U5A cells (non-responsive to type 1 IFNs) PDE12 inhibitors had no anti-viral effect in an EMCV lytic assay indicating that the antiviral properties were dependent on IFN production and not due to direct effects of the drugs on viral replication (data not shown).

Whilst tight regulation of RNAseL activity is required to avoid cell damage, an advantage of targeting PDE12 is that the pathway requires activation by dsRNA thereby providing specificity for those cells which are infected with viruses. Homozygous deletion of PDE12 results in embryonic lethality but heterozygous deletion has no impact on development and animals are phenotypically normal. Higher levels of OAS activity have been reported in rapidly proliferating cells which may explain the effects in early development and might raise the possibility of bone marrow or gastrointestinal side effects if PDE12 inhibition were used for prolonged periods. However, no abnormalities in haematological indices, bone marrow histology or gastrointestinal effects were seen in the rodent toxicity experiments conducted (by Covance) prior to the *in-vivo* antiviral experiments [data not shown].

As dsRNA commonly occur in viral replication cycles, there is broad potential for PDE12 inhibition as an antiviral. However, the OAS-RNAseL pathway is ubiquitous across mammals, birds and reptiles and numerous viruses have developed strategies to resist RNAseL activity. Examples include the influenza virus NS1 protein which binds to the dsRNA binding site on OAS and a conserved region in the RNA of poliovirus binds directly to RNase-L and inhibits substrate cleavage. Members of the coronavirus and rotavirus families encode their own 2’5’ PDE enzymes to regulate RNAseL activity. Our data imply that these resistance mechanisms may be overcome, possibly through inhibition of the viral 2’5’-PDE by the PDE12 inhibitors, although this has not yet been tested. Furthermore, the genetic association of OAS polymorphisms with both HCV and SARS-CoV-2 outcome (4,5) suggest that the pathway is critically important in the innate immune response to these pathogens through the regulation of viral replication.

OAS-RNAseL and PKR pathways are activated in MAVS knockout A549 ACE-2 transfected cells (3) demonstrating that SARS-CoV-2 can induce these host antiviral pathways despite minimal IFN production (3). More recently the importance of this pathway has been demonstrated in SARS-CoV-2 infection by the observation that viral replication is increased by RNAseL knockout (3).

Direct acting antiviral drugs are potent and due to their specificity, less likely to cause side effects. However, the emergence of drug-resistant variants is a constant threat. The mechanisms of action of an OAS-RNAseL-PDE12 inhibitor therapeutic approach, makes it unlikely that viral sequence variation would result in resistance facilitating the potential use of a single agent against a wide range of viral isolates as seen in SARS-CoV-2 infection (Delta and Omicron variants). In addition, viral diagnosis is required before virus specific inhibitors can be deployed in any illness thereby delaying the start of treatment, whereas this is not a requirement for the use of drugs that target host proteins.

In this program we have shown in heterozygote PDE12 KO mice that the PDE12 gene product can be safely reduced in adult mice, although it is essential for foetal development preventing easy development of homozygote KO mice. In PDE12 KO cells (A549 cells with PDE12 deleted using CRISPR/Cas9) we have shown enhanced activity of the 2’*5*’OAS/RNAseL pathway and enhanced anti-viral activity. Finally, the lead drugs are not toxic in initial cellular viability studies and non-toxic at the therapeutic range in rodent studies (Covance data).

## Acknowledgements

Maricela Torres performed *in vitro* and *in vivo* studies at UTMB.

## Financial Disclosure Statement

The work described herein was partly funded by Riotech or investigator own funds.

---

**Supplementary Table 1:** Activity against other phosphodiesterases (PDE12 inhibitor specificity). CO-17 specificity profiling against a panel of phosphodiesterases (PDEs) was outsourced to SB Drug discovery (Glasgow, Scotland, UK).

| **PDE** | **Percentage Inhibition (%)** |  |
| --- | --- | --- |
|  | **CO-17** | **Standard** |
| **PDE1A3** | **27.2** | **59.1** |
| **PDE2A3** | **73.1** | **79.4** |
| **PDE3Cat** | **17.2** | **51.9** |
| **PDE4Cat** | **50.2** | **76.2** |
| **PDE5Cat** | **27.8** | **76.8** |
| **PDE7A1** | **74.9** | **80.3** |
| **PDE8A1** | **6.5** | **41.2** |
| **PDE9A1** | **1.0** | **83.6** |
| **PDE10A1** | **51.3** | **75.1** |
| **PDE11A1** | **39.3** | **47.0** |

## Notes

### Competing Interest Statement

The authors have declared no competing interest.

## References

1. Sadler A J and Williams B R (2008). Interferon inducible antiviral effectors, Nat Rev Immunol 8: 559–568.

2. Kubota K, Nakahara K, Ohtsuko T et al (2004). Identification of 2’ phosphodiesterase which plays a role in the 2’5’ system regulated by interferon. J Biol Chem 279;37832–37841.

3. Li Y; Renner D M; Comas C E et al (2021). SARS-CoV-2 induces double stranded RNA-mediated innate immune responses in respiratory epithelial cells and cardiomyocytes. PNAS 118: e2022643118.

4. Knapp S, Yee LJ, Frodsham AJ, Hennig BJ, Hellier S, Zhang L, Wright M, Chiaramonte M, Graves M, Thomas HC, Hill AV, Thursz MR (2003). Polymorphisms in interferon-induced genes and the outcome of hepatitis C virus infection: roles of MxA, OAS-1 and PKR. Genes Immun 4:411–9.

5. Pairo-Castineira E, Clohisey S, Klaric L, et al. (2021). Genetic mechanisms of critical illness in COVID-19. Nature 591:92–98.

6. Wood E R, Bledsoe R, Chai J et al (2015) The role of PDE12 inhibitors as a negative regulator of the innate immune response and the discovery of antivirals inhibitors. J Biol Chem 290: 19681–96.

7. McAuley A J and Beasley D W (2016). Propagation and titrations of West Nile Virus on Vero Cells. Meth Mol Biol 1435: 19–27.

8. Bewley KR, Coombes NS, Gagnon L, et al. Quantification of SARS-CoV-2 neutralizing antibody by wild-type plaque reduction neutralization, microneutralization and pseudotyped virus neutralization assays. Nat Protoc. 2021 Jun;16(6):3114–3140. doi: 10.1038/s41596-021-00536-y. Epub 2021 Apr 23. PMID: 33893470.

